# Long-read shotgun metagenome sequencing using PromethION uncovers novel bacteriophages, their abundance, and interaction with host bacterial immunity in the oral microbiota

**DOI:** 10.1101/2020.03.13.989996

**Authors:** Koji Yahara, Masato Suzuki, Aki Hirabayashi, Yutaka Suzuki, Yusuke Okazaki

## Abstract

Bacteriophages (phages), or bacterial viruses, are very diverse and highly abundant worldwide, including human microbiomes. Although a few metagenomic studies have focused on oral phages, they relied on short-read sequencing. Here, we conducted a long-read metagenomic study of human saliva for the first time using PromethION that requires a smaller amount of DNA than PacBio. Our analyses, which integrated both PromethION and HiSeq data of >30 Gb per sample, revealed N50 ranging from 187-345 kb and thousands of contigs with >1 kb accounting for > 99% of all contigs on which 94-96% of HiSeq reads were mapped. We identified hundreds of viral contigs (95 phages and 333 prophages on an average per sample); 0-43.8% and 12.5-56.3% of the “most confident” phages and prophages, respectively, didn’t cluster with those reported previously and were identified as novel. Our integrated analyses identified highly abundant oral phages/prophages, including a novel *Streptococcus* phage cluster and nine jumbo phages/prophages. Interestingly, 86% of the phage cluster and 67% of the jumbo phages/prophages contained remote homologs of antimicrobial resistance genes, suggesting their potential role as a source of recombination to generate new resistance genes. Pan-genome analysis of the phages/prophages revealed remarkable diversity, identifying 0.3% and 86.4% of the genes as core and singletons, respectively. Functional annotation revealed that the highest fraction of the core genes was enriched in phage morphogenesis, followed by the fraction enriched in host cellular processes. Furthermore, our study suggested that oral phages present in human saliva are under selective pressure for escaping CRISPR immunity.

**Importance:** Despite the abundance and grave implications oral bacterial viruses in health and disease, little is known regarding the different groups of oral bacterial viruses, their relative abundances under various conditions, and their activities. We provided answers to these questions for the first time utilizing a recently developed sequencer that can capture and sequence long DNA fragments, including viruses, and requires only a small amount of DNA input, making it suitable for analyzing human oral samples. We identified hundreds of viral sequences, including “jumbo” viruses and a distinctive group of highly abundant oral viruses, which often contained parts of antimicrobial resistance genes; the entire repertoire of these viral genes showed remarkable diversity and supported a recently proposed hypothesis that phages modulate oral microbiota through multiple mechanisms. We also revealed genomic signs of coevolution of viruses and host bacteria that have been missed in large viromic studies in humans.

## Introduction

Human microbiomes are of enormous interest to researchers (1, 2) and have been model systems for studying polymicrobial communities (3, 4). Interspecies networks within the microbiome can modulate energy metabolism pathways and affect human health. The two main human microbiomes are intestinal and oral microbiota, which harbor hundreds of coexisting species, including bacteria and viruses. Among them, bacteriophages (phages), or bacterial viruses, which are the most abundant and diverse biological entities in the planet, are present in both intestinal and oral microbiota with implications in health and disease; however, there have been few recent studies on their role in the human microbiome(3, 5, 6).

With the rise of next-generation metagenome sequencing technologies, human gut virome studies have increased rapidly (7). For example, there have been more attempts to characterize “healthy gut phageome” since a study in 2016 conducted deep-sequencing of DNA from virus-like particles (VLP) and revealed the presence of completely assembled phage genomes in 64 healthy individuals around the world (8). Attempts have also been made to explore the associations between human gut virome alterations and diseases (7, 9, 10). However, only a small number of metagenomic studies focused on oral phage communities (5), including a study in 2014 that reported an alteration of virome composition in subjects with periodontal disease (11). The largest oral virome study was conducted in 2015, which generated and analyzed more than 100 Gb shotgun sequencing data from 25 samples (20 dental plaque specimens and 5 salivary) to primarily explore the phage-bacteria interaction network (12). More recently, shotgun metagenome sequencing of 3,042 samples from various environments, including the human oral cavity, was conducted in a project aimed at uncovering the Earth’s virome (13), which achieved an almost 3-fold increase in the metagenome samples (14). Re-analysis of the Earth’s virome data revealed signatures of genetic conflict invoked by the coevolution of phages and host oral bacteria enriched in the human oral cavity (15), suggesting it as an attractive system to study coevolution using metagenomic data. Such metagenomic signs of coevolution have been missed in previous large viromic studies in humans (3).

These previous studies, however, were based on short-read sequencing data generated using Illumina sequencer. However, using contigs assembled from short-reads as short as 2 kb, a popular phage search tool could only identify up to 15% of phage regions, while its sensitivity increases to ∼90% in contigs assembled from reads > 20 kb (16). Therefore, in principle, it is much more desirable to use long-read sequencing data to explore phages and their relationships with host bacteria in metagenomic analyses. Recently, a pioneering study used PacBio sequencing and conducted a long-read metagenomic exploration of extrachromosomal mobile genetic elements, such as phages in the human gut, and generated contigs of 71 plasmids and 11 phages including complete genomes of 5 diverse crAssphages from 12 human fecal samples (17). PacBio sequencing, however, requires relatively high DNA input (∼5µg for standard library protocol) (18), which is usually difficult to obtain from standard amounts of human oral specimens; our preliminary experiment showed that the average amount of DNA extracted from 1 mL saliva of healthy individuals is 2.4 µg, which is smaller than the requirement of PacBio.

In this study, we conducted a long-read shotgun metagenomic study using PromethION, a recently developed high-throughput nanopore sequencer that requires a smaller amount of DNA than PacBio and is thus, revolutionary for oral microbiota research. We also used Illumina HiSeq for sequencing the same samples, enabling error correction and abundance quantification of contigs assembled from the long-reads. Our analyses integrated the deep-sequencing data of PromethION and HiSeq and uncovered hundreds of metagenome-assembled viral genomes, including a highly abundant phage cluster and jumbo phages/prophages (with > 200 kb); the characteristics of these genes and the metagenomic signs of coevolution indicated that the oral phages can evade CRISPR immunity.

## RESULTS

### DNA amount and metagenome sequencing

The concentrations of DNA extracted by the method of Kim, Suda, Hattori et al. (19) from two 1 mL saliva samples taken from four healthy volunteers are shown in Fig. 1A. The range of concentrations varied from 24.5-94.1 ng/μl (average was 63.8 ng/μl), corresponding to at least 1.2 μg total DNA per sample, which meets the input requirement (1.0 μg) of the ligation sequencing kit for PromethION. The amount of metagenome sequencing data obtained for the different samples of each individual using HiSeq and PromethION is shown in Fig. 1B. The range of sequencing data obtained through HiSeq varied from 37.2 Gb to 55.5 Gb for HiSeq (purple in Figure 1B), while it was 38.8 Gb to 90.1 Gb for PromethION (red in Fig. 1B).

**Fig. 1.**
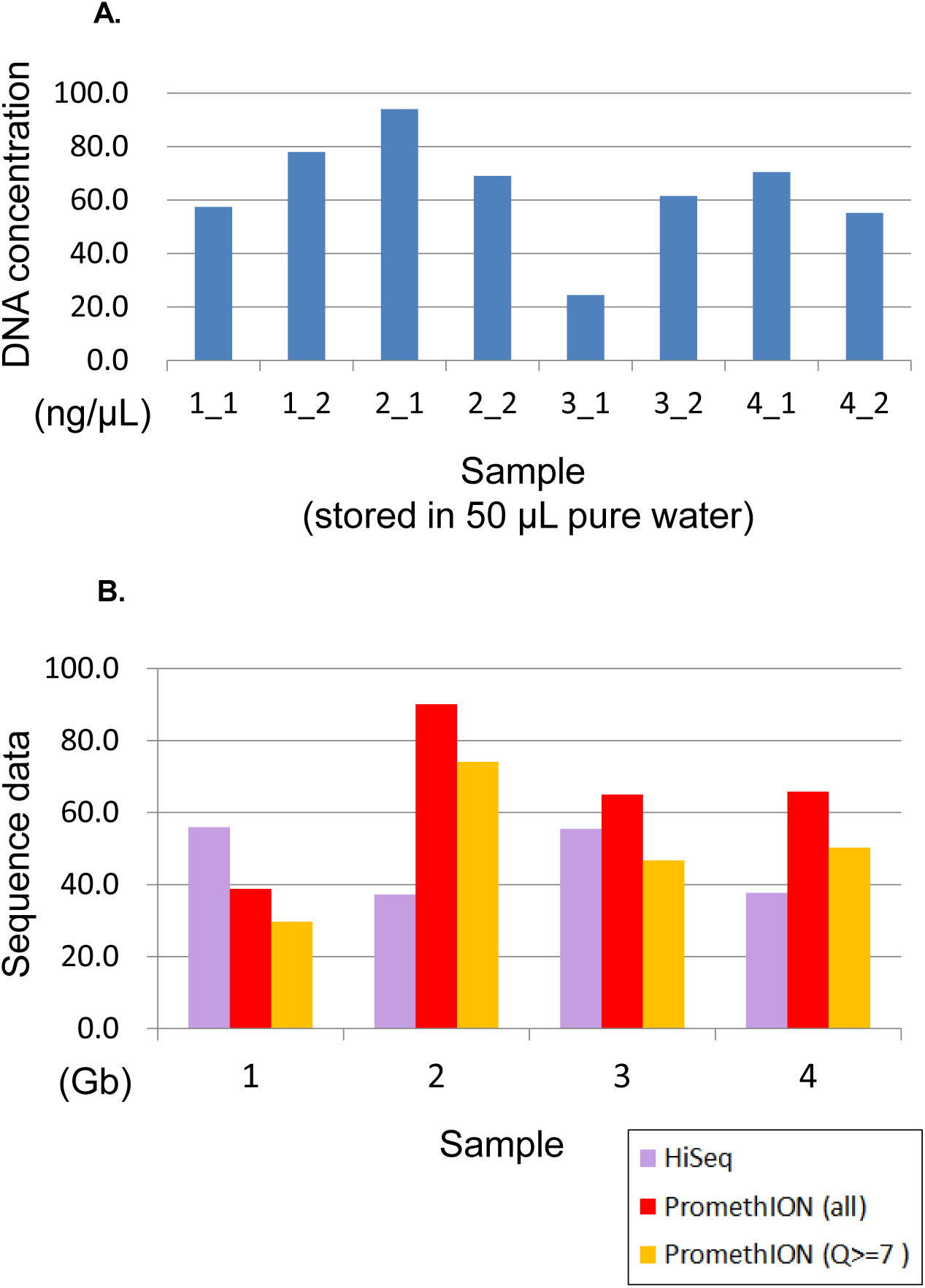
(A) Concentration of DNA extracted from saliva samples. The prefix numbers “2_” and “1_” indicate individual numbers; e.g., “1_1” and “1_2” indicate the 1^st^ and 2^nd^ saliva samples collected from the 1^st^ individual **(B) The amount of generated sequence data.** “Q ≥ 7” (Orange) represents a subset of PromethION reads with average quality score ≥ 7.

### Preprocessing and viral dark matter in human saliva

Preprocessing of the HiSeq data using the EDGE pipeline (20) discarded 0.15-0.22% reads and trimmed 0.31-0.41% of bases by the initial quality control. It then removed 0.20-0.93% of the filtered reads that were mapped to the human genome, which was unexpectedly small; this was likely because of our protocol using a saliva DNA collection kit for microbes and viruses followed by the enzymatic DNA extraction. Preprocessing of the PromethION data discarded 17.9-28.2% of bases (9.1-18.3 Gb) by excluding reads with average quality score < 7 (Fig. 1B), and then removed 0.2-9.7% of the filtered reads that were mapped to the human genome.

Taxonomic profiling using Kraken (21) and its full database for assigning taxonomic labels to the short-reads assigned 0.02-0.2% to “Viruses,” whereas 33.0-55.8% remained unmapped to the database, which is expected to correspond to the viral dark matter manifesting an inability to obtain functional or taxonomic annotations for most of the surveyed sequence space (22). The proportion of unmapped reads was smaller than those (63–93%) reported in marine viral metagenomics (23) but was found to be still dominant in human saliva.

### Length, N50 of contigs assembled from the long-reads

Using a large amount of long-reads (13 kb length on average, Fig. S1), we assembled the long-reads using the recently developed assembler Flye (24) with the “--meta” option followed by error correction based on smapping the HiSeq short-reads to assembled contigs. 94-96% of the HiSeq reads were mapped to the assembled contigs, confirming they well represented viral diversity in the environment. The number of contigs (≥ 1 kb, accounting for > 99% of all contigs in each sample) after the assembly and error correction ranges 2865-5574 per sample with an average of 3802 (Fig. 2A). The N50 ranges from 187 kb to345kb, with an average of 249 kb (Fig. 2B). Nucleotide sequences of all the contigs after the assembly and error correction are downloadable at https://figshare.com/s/e211dd1ab1a77ab94e6f. On the contrary, if we conducted the hybrid assembly implemented in SPAdes (25), the N50 was found to be much smaller (11.8 kb on average, dashed line in Fig. 2B).

**Fig. 2.**
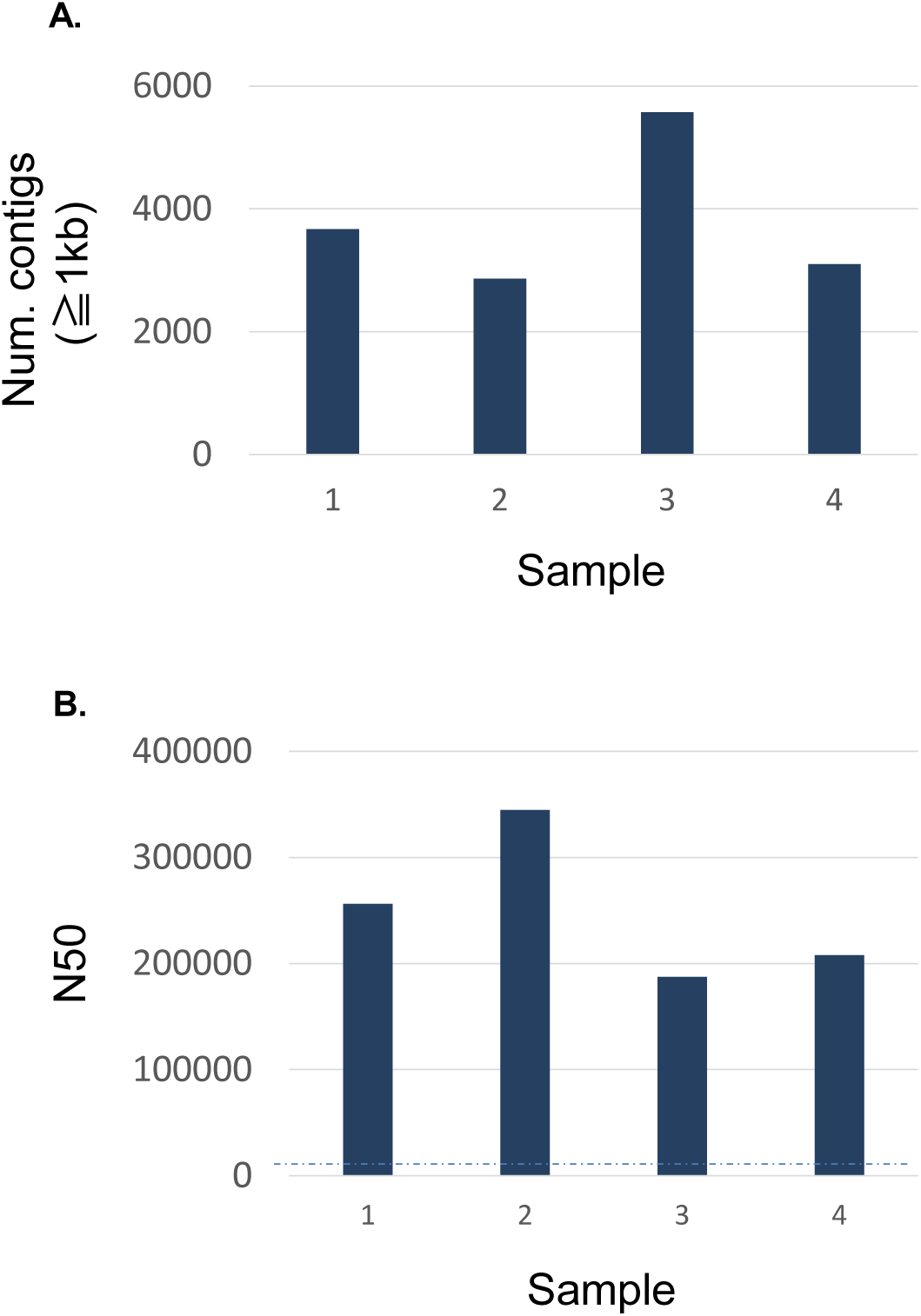
Assembly statistics. (A) The number of contigs with ≥ 1 kb. (B) N50. The dashed line indicates the average of contigs generated by the hybrid assembly rather than the long-read assembly.

### Novel phages/prophages and their taxonomic assignments

For the contigs after the long-read assembly and error correction, we conducted computational classification using VirSorter and identified phage or prophage genome sequences based on the presence of viral ‘hallmark’ genes encoding for components found in many virus particles, together with a reference database of genomes from many viruses (26, 27). The phage or prophage regions identified by VirSorter are classified into category 1 (“most confident”), 2 (“likely”), 3 (“possible”) phages and category 4 (“most confident”), 5 (“likely”), 6 (“possible”) prophages (Fig. S2). The “phages” here include various types of viral sequences outside of the main host chromosome, for example, extrachromosomal prophages (27). The number of each viral category identified in this study is shown in Fig. S2, and the list of all identified viral sequences is shown in Table S1. Because of difficulty to automatically and reliably exclude false positives from the “possible” candidates, we focused on the “most confident” and “likely” phages and prophages (the nucleotide sequences are available at https://figshare.com/s/e211dd1ab1a77ab94e6f). We also applied Contig Annotation Tool (CAT) (28) for taxonomic classification of the contigs based on amino acid sequence searching of each ORF against NCBI nr database followed by a voting approach by summing all scores from ORFs supporting a certain taxonomic classification (superkingdom, phylum, class, order, family, genus, and species, separately) and checking if the summation exceeds a cutoff value (by default 0.5 × summed scores supporting a superkingdom across ORFs). The number of contigs classified as viral by CAT was, however, only ten across the four samples, all of which were included in those by VirSorter.

In order to examine the novelty of the contigs, for each phage or prophage, we conducted genome clustering (29) with oral viral sequences stored in the largest database (IMG/VR v2.0) of cultured and uncultured DNA viruses (30) developed based on the Earth’s virome project (13). Those remaining not clustered with a viral sequence in the database (indicated as empty in “clustered with IMG/VR v2.0” column in Table S1), were considered as novel, and its number and proportion stratified by the “most confident” and “likely” categories are shown in Fig. 3. Among the “most confident” phages and prophages, 0.0-43.8% and 12.5-46.4% were novel, while among the “likely” phages and prophages, 41.7-59.3% and 55.7-73.7% were novel, respectively (colored in yellow in Fig. 3).

**Fig. 3.**
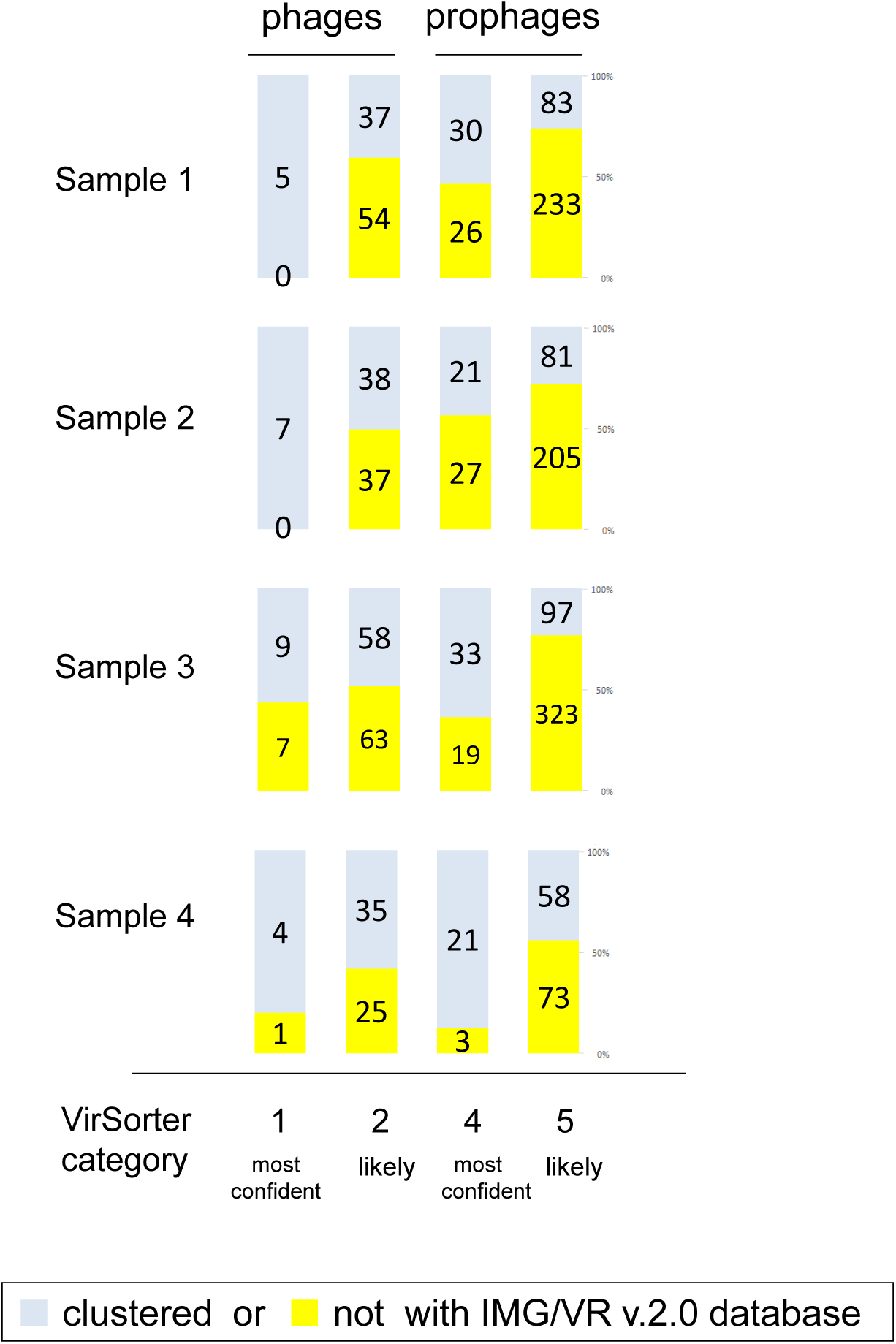
The number and proportion of novel viral sequences identified in each sample and stratified by the “most confident” and “likely” phages and prophages. Yellow: not clustered with viral sequences in the IMG/VR v2.0 database, and regarded as the novel.

Furthermore, for each phage or prophage, we conducted taxonomic assignments based on the extent (network) of gene sharing between a query and the reference viral genomes using vConTACT v2.0 (31). The genus-level assignment was possible only for 3, 1, 1, 3 phages in each of the four samples, respectively. Family-level assignment (indicated in “assigned family” column in Table S1) was possible for 35.9-50.7% of the known phages clustered with the viral sequences in the IMG/VR v2.0 database, and for 38.9-53.8% of the novel phages (Fig. S3). However, the family-level assignment could not be achieved for any of the prophages.

### Highly abundant phages/prophages

The estimated abundance of each viral sequence is shown in Table S1. Rank abundance in each sample (Fig. S4) seems to follow the power distribution as reported in ocean and lake virome studies (32, 33). For top 10% abundant viral sequences in each sample (“top 10% among category 1, 2, 4, 5” in Table S1), we conducted a recently developed gene calling specifically designed for phage genomes (34) that are very compact and often have overlapping adjacent genes (10.5-39.4% in the top 10% abundant viral sequences). The iterative protein searches using the HMM-HMM–based lightning-fast iterative sequence search (HHblits) tool that represents both query and UniProt database sequences by profile hidden Markov models (HMM) for the detection of remote homology (35), annotated 53.5% (11840 out of 22113) genes. After excluding 32 questionable viral sequences (“top 10% among category 1,2,4,5 (questionable)” in Table S1) without carrying any annotated gene related to phage morphogenesis or transposase, we identified 129 predominant phages/prophages across the 4 samples, 61.7% of which were novel because they did not cluster with the viral sequences in the IMG/VR v2.0 database. Self-alignment of each viral contig and manual examination of the dot plot confirmed the integrity and absence of redundancy in their assembly. Nucleotide sequences of the predominant 129 phages/prophages are downloadable at https://figshare.com/s/87a80593aa656ac5567b.

A proteomic tree based on genomic similarity (normalized tBLASTx scores) (Fig. S5) (36) revealed a notable cluster (Fig. S5) that contains 21.7% (28 across the 4 samples, Table S2) of the predominant phages/prophages, which was largest among clusters that can be seen in the tree. Contigs carrying the predominant phages/prophages accounted for 19.0%, 28.8%, 3.4%, and 8.8% abundance (Table S1) among the “most confident” and “likely” viral contigs of each sample. Analysis of the cluster using a larger proteomic tree, including other reference viral sequences, revealed its location distinctively in a *Siphoviridae* and among *Streptococcus* phages (Fig. 4). Examination of their genes revealed that 86% (24 out of the 28) of them encoded remote homologs of antimicrobial resistance genes with > 99% estimated probability to be (at least partly) homologous to the gene sequences (“CDS related to resistance” Table S2). Genomic context and coverage of HiSeq read of one of the predominant prophages is shown in Fig. 5, in which three remote homologs of antimicrobial resistance genes are colored in red. The remote homologs of genes for phage resistance proteins and acid resistance proteins were also found in 7% and 11%, respectively.

**Fig. 4.**
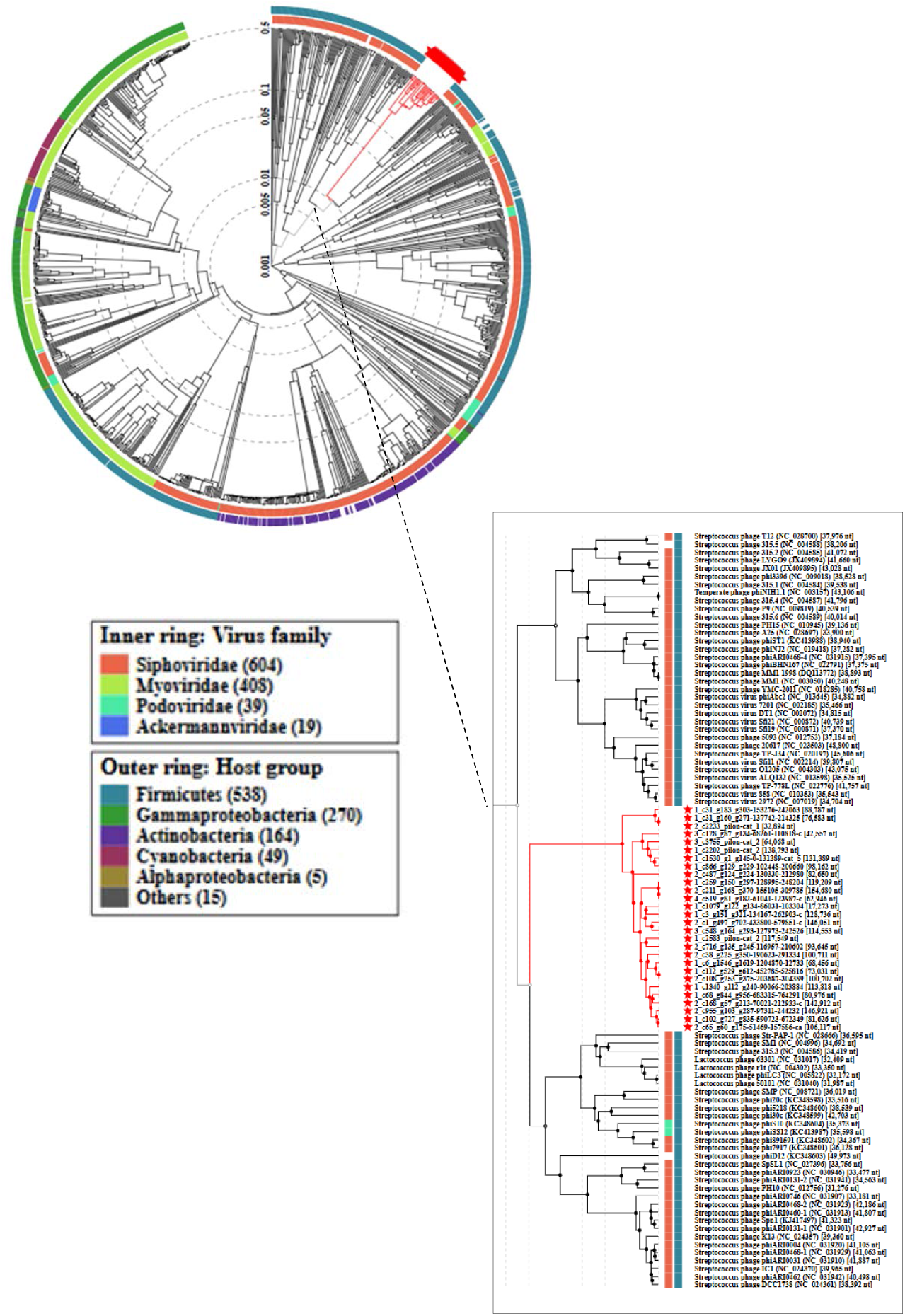
Proteomic tree showing a cluster of highly abundant oral phages. The red lines at the top right and bottom right indicates the cluster consisting of the 28 abundant phages found across the 4 samples and listed in Table S2.

**Fig. 5.**
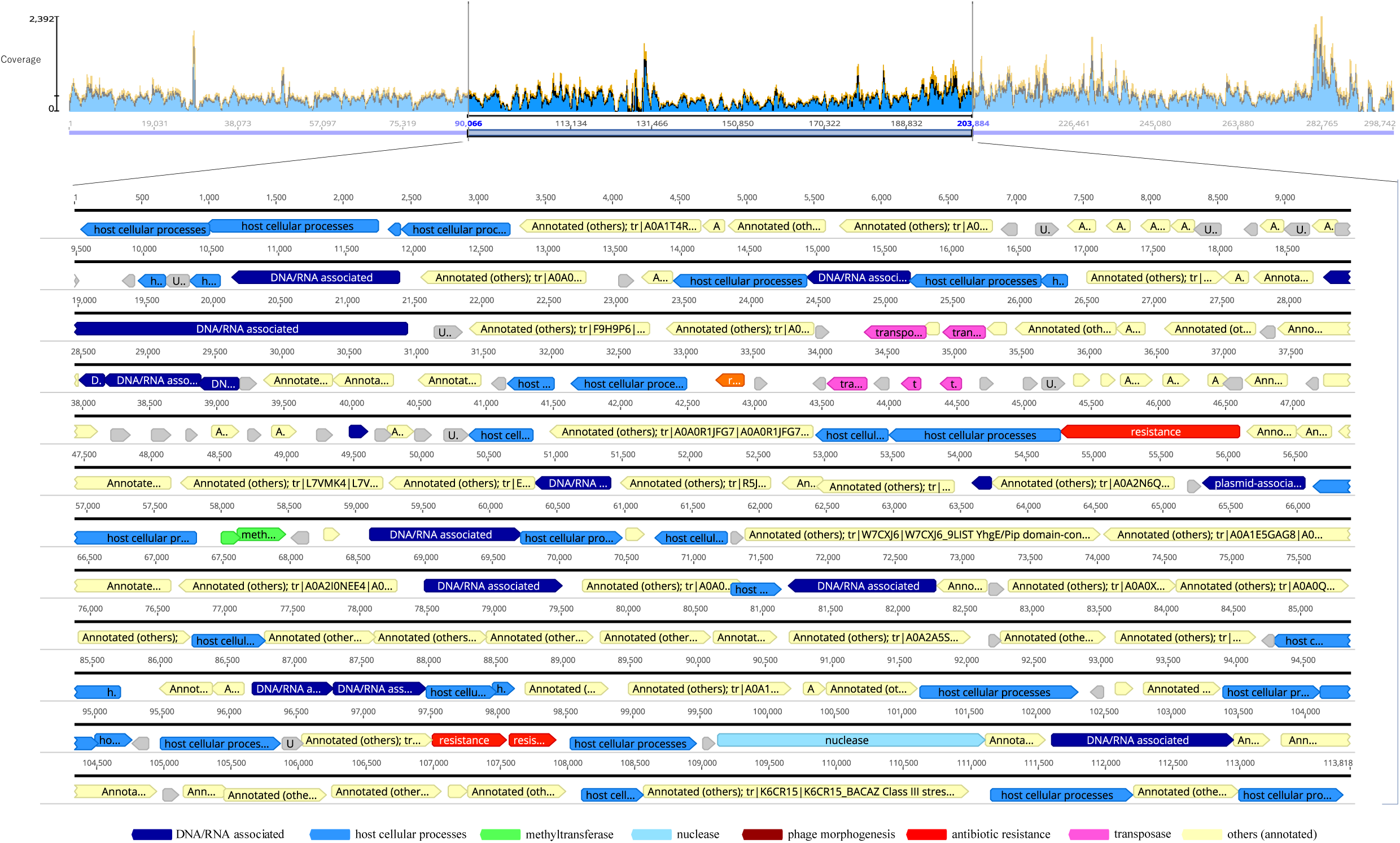
A highly abundant oral prophage. Top: coverage plot based on mapping short-reads against it. Bottom: Genes characterized by the HMM-based iterative protein searches are shown with colors according to their functional categories.

### Jumbo phages/prophages

The distribution of the size of the phages and prophages is shown in Fig. S6. The average, median, and interquartile range (IQR) were 57.9 kb, 44.4 kb, 33.9-76.9 kb for the phages, respectively. For the first time in oral microbiota publications, we discovered six phages with genomes larger than 200 kb (Fig. S6, Table 1), that were classified as jumbo, and had been rarely found but recently began to be identified across Earth’s ecosystems (37, 38). We also discovered five jumbo (>200 kb) prophages. Nucleotide sequences of the jumbo phages and prophages are downloadable at https://figshare.com/s/95f9c3cbb074b2782ccf. Manual examination based on self-alignment and dot plots revealed one of them (contig_659 of the 3^rd^ sample) contained a redundant sequence region spanning approximately 160 kb that might be resulted from an assembly error. After excluding it, 90% of them (nine out of the ten) had at least one phage hallmark gene. One of the remaining contigs with 223995 bp without a phage hallmark gene (contig_811 in the 2^nd^ sample) was classified as a phage by VirSorter, but 8.9 kb region in the middle (94241-104074) encodes three remote homologs of Type IV secretory system protein for conjugative DNA transfer as well as that of plasmid segregation protein. These results suggested it is rather a plasmid-like element as reported previously (39).

**Table 1.**
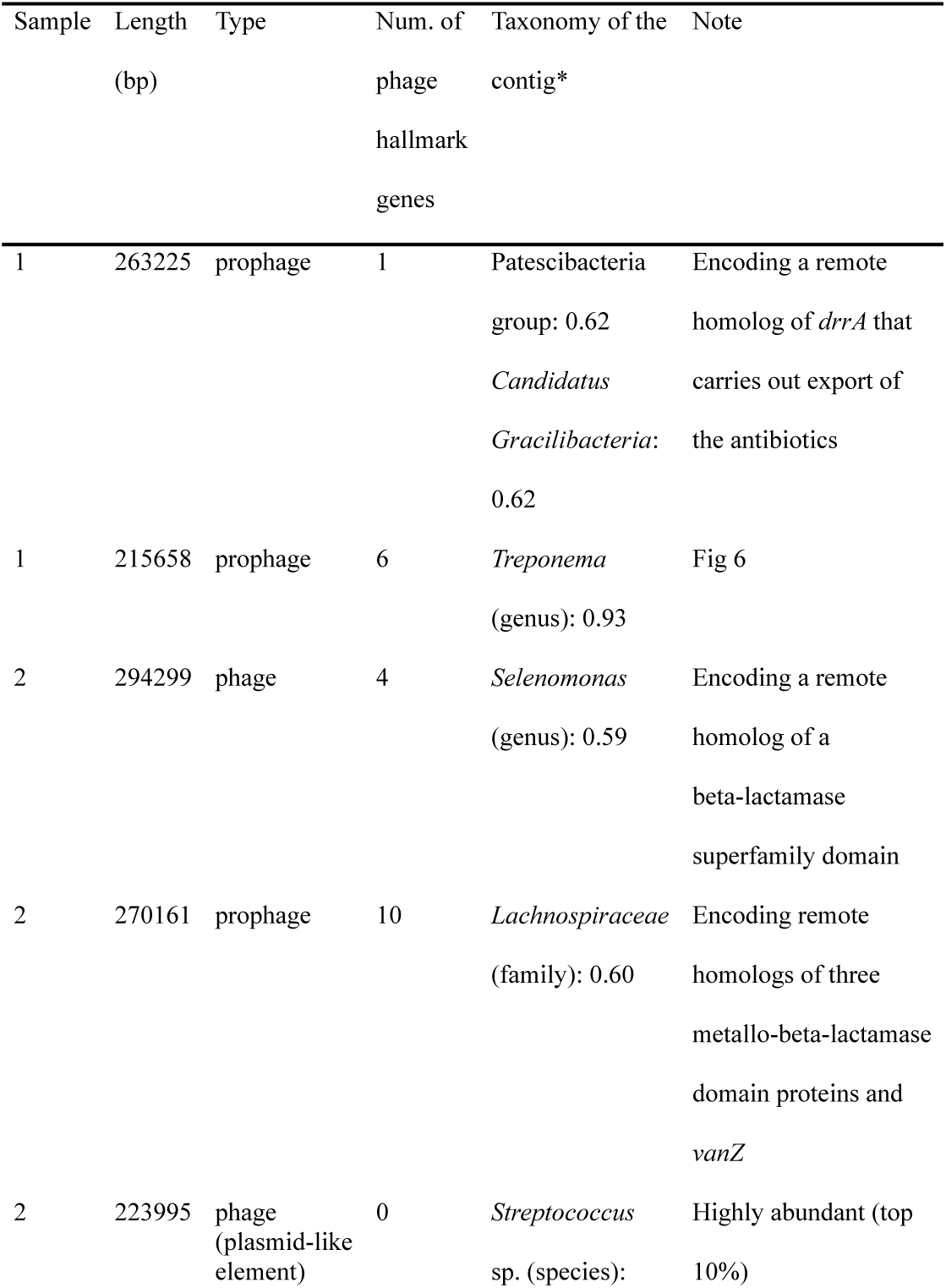
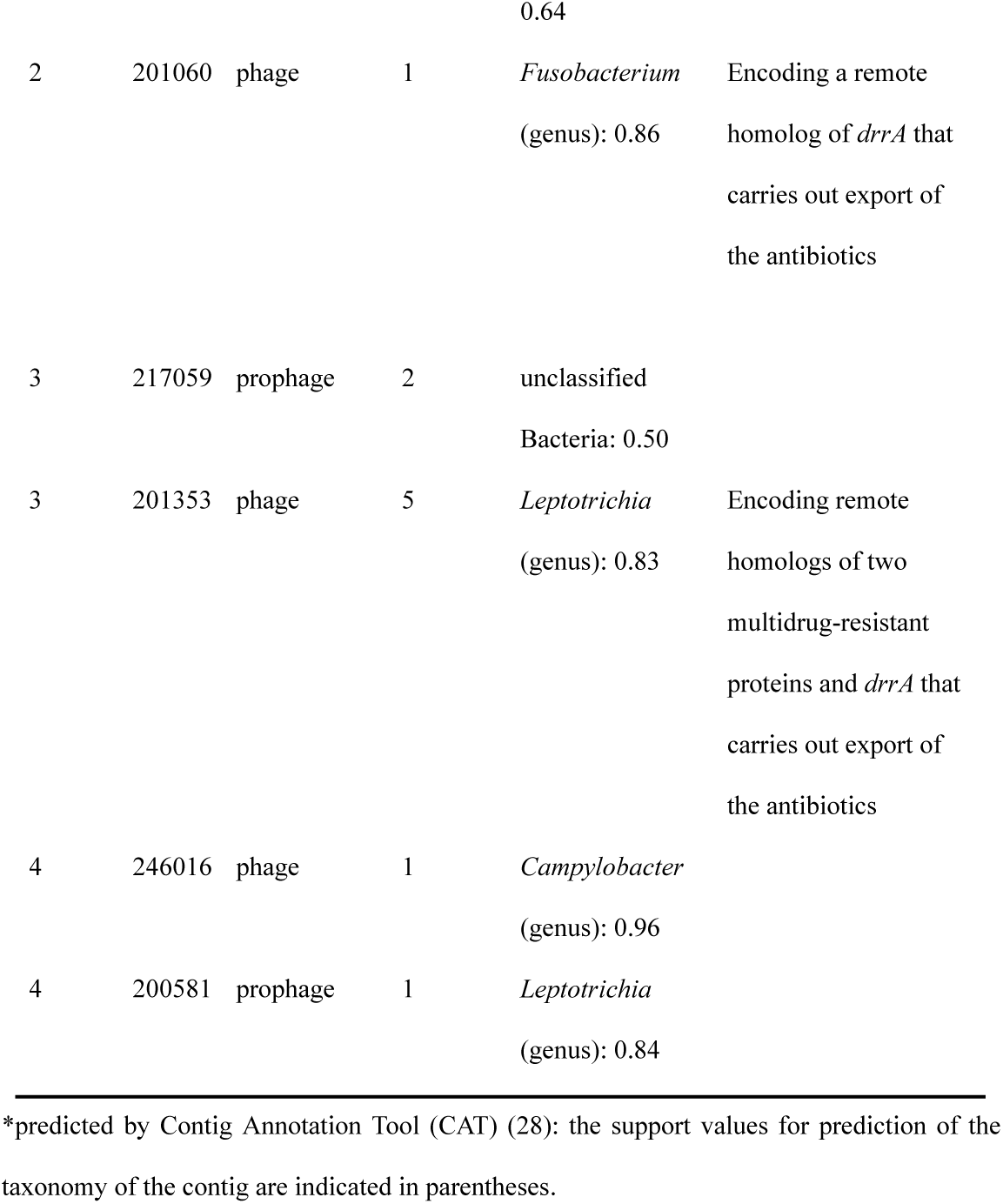
Detected jumbo phages and prophages.

For the jumbo phages and prophages, as explained above, we conducted the gene calling specifically designed for phage genomes (34) and found that 13.2-32.6% of the genes were annotated. Regarding the annotated genes, similar to the highly abundant oral phages/prophages examined above, remote homologs of antimicrobial resistance genes (“Note” in Table 1) were found in 67% (six out of the nine) of the phages/prophages carrying the phage hallmark genes. An example of jumbo prophage embedded in a bacterial chromosome is shown in Fig. 6, in which seven remote homologs of antimicrobial resistance genes with > 99% estimated probability are located (reds in Fig. 6). Six out of the seven were remote homologs of beta-lactamases and formed a cluster in a genomic region (approximately 15.5 kb-17.7 kb; Fig. 6). 31-75 amino acid residues were in the pairwise HMM alignment between the query and beta-lactamases in the UniProt database. The seventh is a remote homolog of *drrA* (77 amino acid residues were in the pairwise HMM alignment) that carries out the export of the antibiotics (40), which was also found in the other three jumbo phages/prophages (Table 1). The taxonomy of the contig was predicted to be the *Treponema* genus, consisting of dozens of species in the human oral microbiota (41). Predicted taxonomy of another contig containing a jumbo prophage was *Patescibacteria*, the recently proposed Candidate Phyla Radiation (CPR) lineage that encompasses mostly unculturable bacterial taxa with relatively small genome sizes (42), including ubiquitous members of the human oral microbiota (43). Overall, there was no overlap in the predicted taxonomies among the jumbo phages/prophages carrying the phage hallmark genes, suggesting they are not confined to specific phylogenetic groups in the human oral microbiota.

**Fig. 6.**
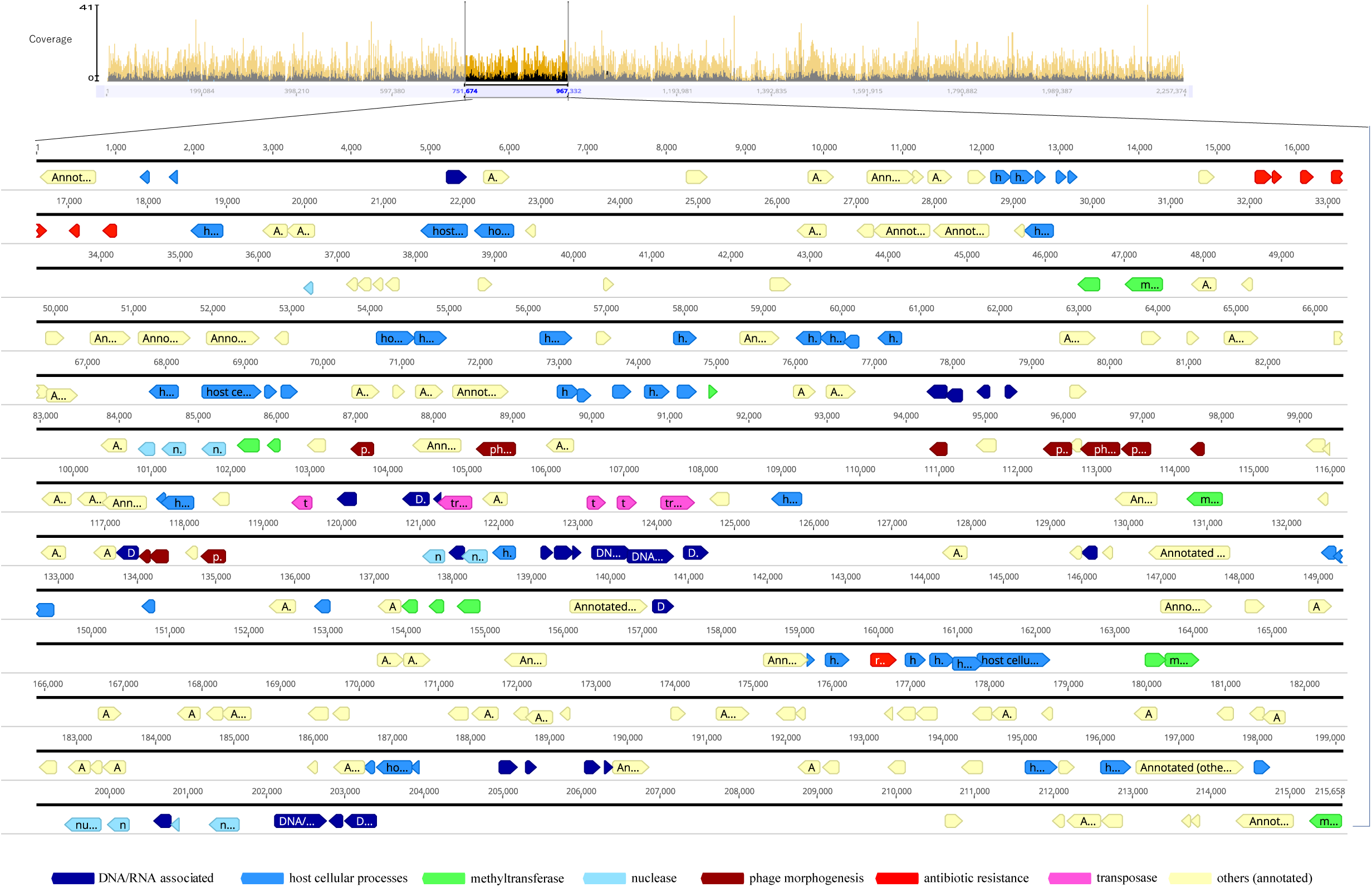
A jumbo prophage. This Fig. is similar to Fig. 5. The prophage region is indicated as a rectangle in the coverage plot spanning the entire contig.

### Characteristics of genes encoded in oral phages/prophages

The pan-genome analysis of all the identified viral sequences to create a gene presence or absence matrix revealed remarkable diversity. Among 115621 different genes (i.e., the number of rows in the matrix), only 0.3% (309) genes were “core” and present (based on 70% amino acid sequence identity by BLASTp) in all the four samples, while 86.4% (998641) genes were singletons (Fig. 7). The amino acid sequence alignments of the 309 core genes are downloadable at https://figshare.com/s/9f76b265f23e23d1e63f. As much as 94.8% (293 out of 309) of the core genes were annotated as hypothetical by a standard annotation program Prokka compatible with the pan-genome analysis (44). We then conducted the iterative protein searches using the HHblits tool that detected remote homologs with > 99% estimated probability for 301 out of the 309 (Table S3). A breakdown of their functional categories (Fig. 7) shows 38.9% were homologs of uncharacterized proteins (gray), while the two most dominant annotated functional categories are phage morphogenesis (25.9%, red) and host cellular processes (14.6%, green) such as transcriptional regulators. Of the remaining 20.6%, DNA/RNA metabolism (6.3%, yellow) and phage lysis (3.3%, light purple) accounted for the half, while DNA recombination (light yellow), virulence (purple), nuclease (blue), methyltransferase (orange), prophage antirepressor (brown), transposase (dark gray), and host”s phage susceptibility (khaki) accounted for the remaining half.

**Fig. 7.**
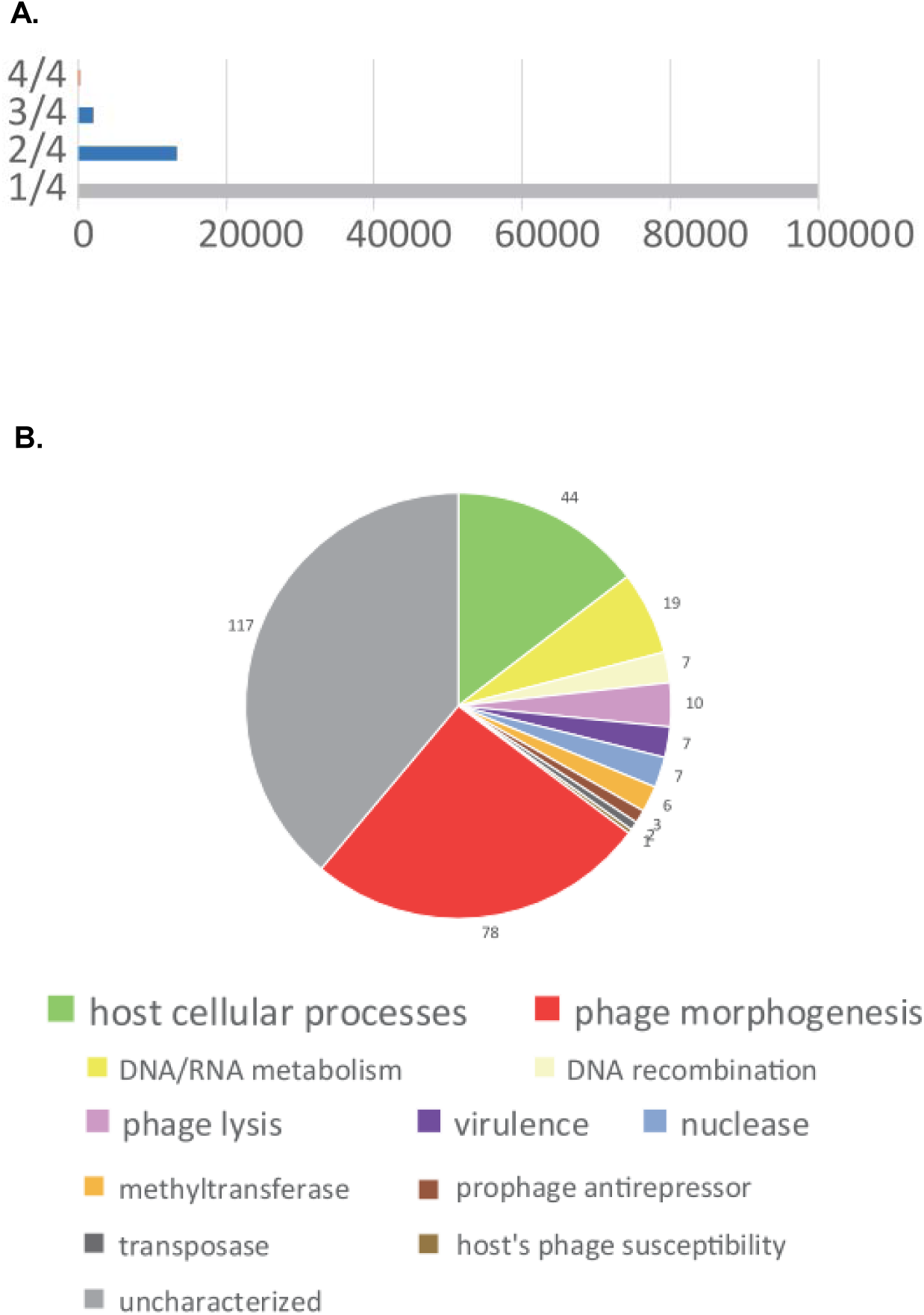
(A) The number of the gene present in 4, 3, 2, 1 out of the four samples. The core (4/4) genes are colored in orange, while singletons are colored in grey. **(B) Breakdown of the core genes.** The number of genes in each category is shown in the pie chart. More detailed information on each gene is shown in Table S3.

### Distribution of CRISPR spacers and its implication to coevolution between oral phages and bacteria

Finally, for all contigs, we detected CRISPR arrays and spacer sequences. In total, we detected 16187 unique spacers across the four samples. The number of spacer sequences counted separately in bacterial contigs and phages/prophages (excluding the “possible” sequences) is shown in Fig. 8A. The average number of spacers in bacterial contigs and prophage sequences was 2503 and 668, respectively (Fig. 8A). Whereas, the spacers in CRISPR arrays were also found in the phage sequences, as reported in other phages (45): 16 spacers on an average in three samples except for the 4^th^ sample (Fig. 8A).. The BLAST search of the spacers in CRISPR arrays located in the phage sequences against two datasets of oral phage/prophage sequences (either in IMG/VR v2.0 database or all the assembled viral contigs in its own sample including the category 3 and 6 (“possible”) candidates), revealed almost no homologous sequences (“protospacers”) (Fig. 8B). The number was only one or five in the 2^nd^ sample and zero in the other three samples. Whereas, the BLAST search of the spacers in CRISPR arrays in the bacterial contigs and prophage sequences against the IMG/VR v2.0 database revealed the on an average 22.2% and 20.3% of the spacers had homologous sequences (“protospacers”) in the database, respectively. In contrast, when the BLAST search was conducted against all the assembled viral contigs in each sample, the average proportion decreased to 1.8% for spacers both in the bacterial contigs and in the prophage sequences. The difference between the two conditions was statistically highly significant (*p* < 10^−15^, chi-square test). Further examination of the result of BLAST search of spacers against IMG/VR v2.0 database and that of the genome clustering conducted above (Fig. 3) revealed that among the oral viral contigs in IMG/VR v2.0 database carrying the protospacers, only 5.2% showed nucleotide similarity enough to be clustered with those identified in the present study, which was significantly lower (*p* < 10^−15^, chi-square test) than the overall proportion of clustering between the two datasets of viral contigs (36.0%, Fig. 3). In other words, most of the oral viral contigs in the IMG/VR v2.0 database carrying the protospacers were outside those identified in the present study.

**Fig. 8.**
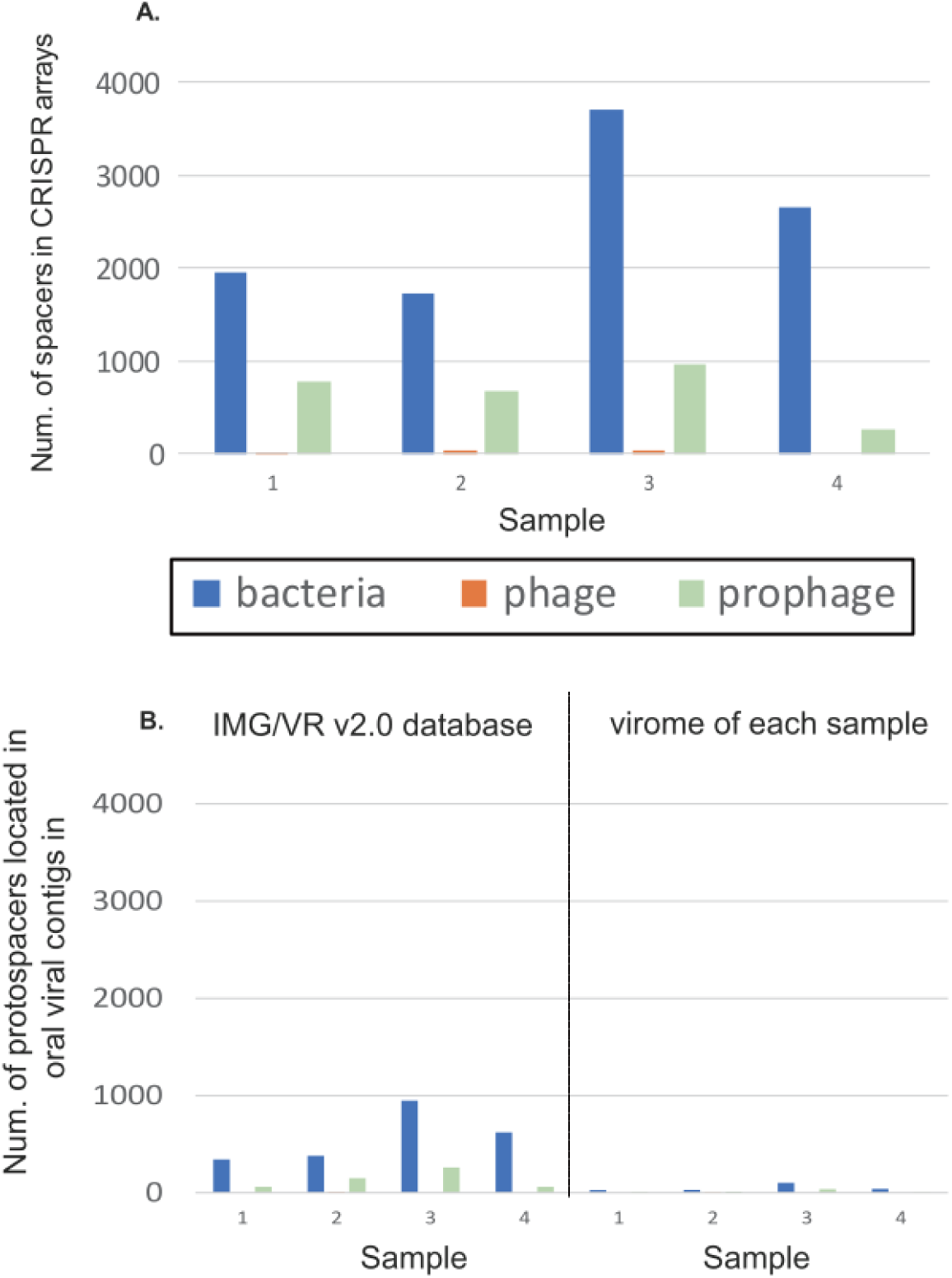
(A) The number of spacers in CRISPR arrays found in the bacterial contig, phage, and prophage regions. (B) The number of protospacers located in oral viral contigs in either IMG/VR v2.0 database (left) or those assembled in each sample (right).

## Discussion

The present study, for the first time, analyzed as much as > 30 Gb metagenomic data of both PromethION and HiSeq per sample. The analysis revealed 40.0-56.3% of the phages and 49.0-72.5% of the prophages were novel (Fig. 3), and identified as much as ten jumbo oral phages/prophages (including the plasmid-like element) in an oral environment, significantly increasing our knowledge about “who is there” in the human oral cavity. Such jumbo phages were previously found among approximately 20 host bacterial genus (38, 46, 47) and recently began to be identified across Earth’s ecosystems (37). However, so far, there is no publication of those in the human oral cavity. Long-read sequencing using PromethION will boost the discovery of such novel jumbo phages/prophage more in various environments.

The analysis also enabled quantification of the abundance of each assembled viral sequence (Table S1), and for the first time, identified the highly abundant oral phages/prophages, some of which formed the novel distinctive *Streptococcus* phage cluster (Fig. 4). There has been no study that demonstrated the abundance of oral phages and quantitatively answered what kind of phages are not only present but also abundant in the human oral cavity. In the *Streptococcus* phage cluster, there was a “most confident” phage (contig_2233 in the 2^nd^ sample) that was similar to a known *Streptococcus* phage SM1 (NC_004996 in Fig. S7) and clustered with 7 viral sequences in the IMG/VR 2.0 database (Table S1). Whereas, 85.7% (24 out of the 28) seemed to be novel as they did not cluster. The genome clustering used cutoffs for obtaining pairwise hits, which are more stringent than the normalized tBLASTx scores used for constructing the proteomic tree. 87.5% of them (21 out of the 24) were “likely” prophages (Fig. S5), of which 2 had genome synteny to the “most confident” phage in the cluster (Fig. S7). Regions outside the synteny of the “likely” prophages did not encode a gene for phage morphogenesis, and might not be viral but host bacterial sequences at both ends mistakenly extracted by VirSorter, making the “likely” prophages not clustered with viral sequences in the IMG/VR 2.0 database. We should keep in mind if that is the case, the proportion of novel “likely” prophages, as well as of jumbo prophages, could be overestimated.

A notable feature in oral microbiota revealed from the largest deep-sequencing data of both PromethION and HiSeq was high frequency of the remote homologs of antimicrobial resistance genes; 67% of the jumbo phages/prophages and 86% of the predominant *Streptococcus* phages cluster. There has been controversy as to how prevalent the antimicrobial resistance genes are really in phages, given that the genes are only rarely directly encoded in publicly available phage genome sequences (48). The remote homologs formed the pairwise HMM alignment not entirely but partly with the antimicrobial resistance genes, and are perhaps unfunctional. However, they might play a role as a source of recombination or horizontal gene transfer to generate a new antimicrobial resistance gene sequence. In the human oral cavity, it is known that the “mosaic” penicillin-resistant genes are generated by recombination among oral streptococci (49) and that the massive diversity of organisms and a large amount of extracellular DNA in oral biofilm matrices are expected to create opportunities of recombination or horizontal gene transfer (4).

Regarding the analysis of the distribution of CRISPR spacers and protospacers, a previous study reported that protospacers were detectable as a small fraction (from 1% to 19%, on average ∼7%) of the spacers (50). The proportion in our study obtained by the BLAST search of the spacers against the oral viral sequences in the IMG/VR v2.0 database was slightly higher than the upper limit (19%). A more recent study of the human gut microbiome (51) reported that the proportion was 23.5% when IMG/VR v2.0 database was used, which is comparable to the findings of our study. When the BLAST search was instead conducted against all the assembled viral contigs in each sample, the proportion was significantly decreased. The possible explanation for this reduction that the assembled viral contigs account for only a small part of the overall oral virome is nullified because 94-96% of the HiSeq reads were mapped to them. However, it suggests that the oral phages currently in human saliva could be under selective pressure escaping from CRISPR immunity, given hundreds of thousands of CRISPR spacer groups are transcribed in the human oral cavity (52). Previously, it was reported that oral streptococcal CRISPR spacers and viruses carrying protospacers coexisted in human saliva of a subject, and CRISPRs in some subjects were just as likely to match viral sequences from other subjects as they were to match viruses from the same subject (53). Compared to the previous study that examined 3473 unique spacers confined to streptococci of 4 subjects (53), we expanded the number to > 4-fold across various bacterial species. Therefore, the result presented in our study is probably more general and reliable than the previous study in understanding the overall relationship between CRISPR spacers and protospacers in the oral microbiota. A recent review also pointed out the possibility of genetic conflict invoked by the coevolution of oral phages and host bacteria using classic Red Queen dynamics (4). The possibility was recently suggested in another study that revealed signatures of highly elevated recombination among oral phage genomes, potentially reflecting the evolutionary conflict in which phages continuously change their genomic sequences by recombination to escape cleavage by the host bacterial immune system in the human oral cavity (15).

Recently, a framework of viral, long-read metagenomics via MinION was proposed, and one of its main advantages was to capture more and longer genomic islands (54). The genomic island was defined in a previous study by coverage of less than 20% of the median coverage of the entire contig and >500 bp in length (55). However, such a decrease in coverage can result from a long read-assembled contig that contains too erroneous regions for correction by short-read mapping. In particular, long-read assembly of low coverage contig is challenging, and detection of a genomic island in such a contig was not practical. This issue is worth to be addressed in future works by exploiting the improvement of per-base accuracy and per-cost throughput of the long-read sequencing platforms.

A remarkable feature we found in the pan-genome analysis was the high diversity of genes encoded by oral phages/prophages: as much as 86.4% of the genes were singletons (i.e., specific to each sample). It is consistent with previous studies of 16S rRNA-based metagenomics reporting individuals’ oral microbiota are highly specific at the species level. Our study further demonstrated the specificity at the gene level based on the very deep shotgun metagenome sequencing.

Furthermore, we conducted identification and HMM-based computational characterization of the core genes of oral phages (most of which were initially annotated as hypothetical) to deepen our understanding of what they are doing. The high fraction of the core genes for host cellular processes after phage morphogenesis supports a recently proposed notion that phages modulate the oral microbiome through multiple mechanisms and represent an additional level of balance required for eubiosis (4). Experimental characterization of the genes in the future will further deepen our understanding of their functions and roles.

In summary, our study demonstrates, for the first time, the power of long-read metagenomics utilizing PromethION in uncovering novel and abundant bacteriophages, characteristics of their genes, and their interaction with host bacterial immunity. Our study will provide a solid basis for utilizing PromethION to study bacteriophages and host bacteria simultaneously, and further explore the viral dark matter in various environments.

## Materials and Methods

### Saliva collection, DNA extraction, library preparation, and metagenome sequencing

Two samples of 1 mL saliva were successively collected and stored using a kit specialized for microbial and viral DNA/RNA (OMNIgene ORAL OM-501) from each of four healthy volunteers (2 men and 2 women aged 35 to 65 years old). From the four individuals, two saliva samples for each were taken and subjected to DNA extraction using an enzymatic method (19). The extracted DNA samples were stored in 50 μL pure water. From the paired samples per person, we used one with a smaller amount of extracted DNA for Nextera XT library construction and genome sequencing with the Illumina HiSeq 2 × 150 bp paired-end run protocol, and another for library construction using the ligation sequencing kit (SQK-LSK109) and PromethION sequencing.

### Preprocessing and initial taxonomic profiling

We used EDGE pipeline version 1.5 (20) for preprocessing (trimming or filtering out reads, and removal of reads mapped to the human genome) of the HiSeq data. For the PromethION data, we used MinIONQC (56) to check diagnostic plots and NanoFilt (57) (“-q 6 --headcrop 75” option) to filter out reads with average quality less than 7 and trim 75 nucleotides from the beginning of a read. We then used Minimap2 (58) to find and remove PromethION reads mapped to the human genome. Initial taxonomic profiling using HiSeq reads was conducted using Kraken (21) and its full database created in March 2017 following https://github.com/mw55309/Kraken_db_install_scripts as a part of the EDGE pipeline. We executed it implemented in metaWRAP (59) as a module that can directly subsample the same number (approximately 120 million) of reads in each sample

### Assembly, error correction, and quality assessment

We used Flye (24) with the “--meta and –genome-size 200 m (a value much higher than known bacterial genome size) “option to assemble the preprocessed PromethION long-reads, followed by mapping the preprocessed HiSeq short-reads to assembled contigs using bowtie2 (60) with “—very-sensitive” option. We then conducted an error correction based on the mapping using a single run of Pilon for each assembled contig (61). It took several days from the mapping to error correction. We also conducted a hybrid assembly implemented in SPAdes with “—meta --nanopore” option (25). Quality assessment for the assembled contigs was conducted using MetaQUAST (62).

### Viral sequence identification, taxonomic classification, abundance quantification, and clustering of assembled contigs

We used VirSorter (26) with the “-db 2” option using the “Viromes” database. We also used CAT (28) with the default database and taxonomy information created in January 2019 and included in the package. Distribution of length of viral sequences identified by VirSorter was examined using JMP Pro version 13 (SAS Institute, Cary, NC, USA). The abundance of each viral sequence was estimated by FastViromeExplorer (63) using randomly selected 123519885 read-pairs (equal to the number of the least sequenced sample) from each of the four samples. A proteomic tree of highly abundant viral sequences (top 10% in each sample) was constructed using ViPTreeGen, and a larger proteomic tree, including the highly abundant viral sequence cluster and other reference viral sequences, was constructed using ViPTree (36) in which their genome alignment was also visually examined. The viral sequences were clustered with oral viral sequences registered in IMG/VR v2.0 database (30) according to a “Viral genome clustering” procedure included in a nontargeted virus sequence discovery pipeline (from step 8 to 11 in (29), using cutoffs of nucleotide sequence similarity ≥ 90%, covered length ≥ 75%, and covered length requiring at least one contig of > 1,000-bp length. Taxonomic assignment of the contigs was conducted using vConTACT v.2.0 (31) with “--db ‘ProkaryoticViralRefSeq85-Merged’ --pcs-mode MCL --vcs-mode ClusterONE” option.

### Functional annotation and pan-genome analysis of viral sequences

The prediction of protein-coding genes in the phages/prophages was conducted using PHANOTATE (34) implemented in multiPhATE (64). For each predicted gene, we conducted iterative protein searches using HHblits (35), which represents both query and database sequences by profile hidden Markov models (i.e., condensed representation of multiple sequence alignments specifying, for each sequence position, the probability of observing each of the 20 amino acids) instead of single sequences for the detection of remote homology. We used the clustered uniprot20_2016_02 database (http://www.user.gwdg.de/~compbiol/data/hhsuite/databases/hhsuite_dbs/), which covers essentially all of the sequence universe by clustering the UniProt database (65) from EBI/SIB/PIR and the non-redundant (nr) database from the NCBI. For all hits with > 99% probability of being true positives, we visualized their genomic locations using Geneious software (Biomatters Ltd., Auckland, New Zealand) and individually examined each annotation. We did not include uncharacterized genes in the visualization because the number was too many for the software. Furthermore, we conducted pan-genome analyses using the Roary pipeline with the “-i 70 –group_limit 500000” option (66) after the prediction of protein-coding genes for every contig using Prokka software with the “–kingdom Viruses” option (44). For the core genes, we also conducted the iterative protein searches using HHblits, individually examined each annotation, and manually made the functional categorization.

### Analyses of distribution of CRISPR spacers

CRISPR arrays were predicted on all contigs using a command-line version of the program CRISPRDetect (67). For each sample, spacers were extracted from the output files and searched using the BLASTn-short function from the BLAST+ package (68) against either oral viral contigs in IMG/VR v2.0 database or those assembled in this study in each sample. The cutoffs were set with at least 95% identity over the whole spacer length and allowing only 1–2 SNPs at the 5’ end of the sequence, according to a procedure in the Earth’s virome project (13).

### Ethics

This study was approved by the ethics committees of National Institute of Infectious Diseases (approval number 931).

### Data availability

The data of PromethION and HiSeq after the quality control and removal of human reads were deposited at DDBJ (with JGA accession number JGAS00000000186) and will be mirrored at NCBI under BioProject accession PRJDB9452.

## Acknowledgments

The computational calculations were done at the Human Genome Center at the Institute of Medical Science (the University of Tokyo). This work was supported by Grants-in-Aid for Scientific Research from the Ministry of Education, Culture, Sports, Science and Technology (MEXT) (19H04846 to K.Y.) and JSPS KAKENHI Grant Number 16H06279 (PAGS). We thank Yutaka Suzuki laboratory members for PromethION sequencing. We thank Ryan Wick and Nick Loman for the instruction of the long-read assembly and thank Matthew B Sullivan, Masahira Hattori, Yosuke Nishimura, Tetsuya Hayashi, Takuro Nunoura, Yoshihiro Takagi, Masaki Shintani and So Nakagawa for discussions.

